# Classification and Evolution of the Plant 4CL/ACS Superfamily and Transcript Analysis in Andean Potato

**DOI:** 10.1101/2021.07.06.451337

**Authors:** Matías Ariel Valiñas, María Luciana Lanteri, Adriana Balbina Andreu, Arjen ten Have

## Abstract

Hydroxycinnamates, intermediates of phenylpropanoid metabolism, can be converted by Class I and Class II 4-coumarate:CoA ligases (4CL) to their activated CoA esters. Class I enzymes are involved in suberin/lignin deposition, whereas Class II enzymes feed flavonoid biosynthesis. Before initiating transcript analysis in Andean potato tubers, we undertook a thorough classification of the 4CL/ACS superfamily to settle a confusing and inconsistent description of the family in literature. Phylogenetic analysis and clustering of sequences from SwissProt and 15 plant species resolved the cytosolic Class I 4CL, Class II 4CL and ACOS5 subfamilies, as well as the peroxisomal subfamilies of Class III 4CL, OPC-8:0-CoA ligase (OPCL) and Uncharacterised ACS I, II and III. Analysis shows that the ACOS5 family, which has oleate as substrate, has a rather different binding pocket. Class III enzymes, which feed ubiquinone biosynthesis, and OPCL enzymes, involved in jasmonic acid biosynthesis, have binding pockets similar to those of cytosolic 4CL enzymes. Differences between Class I and II suggest that CoA governs flux regulation, and that hydroxycinnamate substrate specificity likely did not constrain their evolution. Notably, using sequences from 200 additional plant species, we show most eudicots possess two and most Poaceae three functionally non-redundant Class I homologues, suggesting that substrate specificity did constrain the evolution of the Class I subfamilies. Finally, we show that Class I and Class II transcript regulation corresponds to the proposed classification and may reflect Class I diversification constrained by different requirements for suberin and lignin deposition in native and wounded potato tuber periderm.

## Introduction

Potato (*Solanum tuberosum* L.) is an important worldwide staple food crop and considered a cheap source of high-quality proteins, minerals, and antioxidants including vitamin C, carotenoids and phenolic compounds (Camire et al. 2009). The biosynthesis of phenolic compounds depends on the phenylpropanoid metabolism, which is considered an important intermediate between primary and specialised metabolism. Previously, we reported on its main component chlorogenic acid (CGA) in relation to the production of suberin (Valiñas et al. 2015), anthocyanins, and flavan-3-ols (Valiñas et al. 2017). We have shown that CGA and anthocyanin appear to be positively correlated in Andean potato tubers. Suberin, CGA and flavonoid production require 4-coumarate:CoA ligase (4CL) enzymes, which use a variety of substrates known as hydroxycinnamates. Like many other enzymes in the phenylpropanoid metabolism, 4CL enzymes show high substrate permissiveness. Figure 1 summarizes the phenylpropanoid metabolism with emphasis towards 4CL enzymes and the central metabolite CGA.

**Fig 1.**
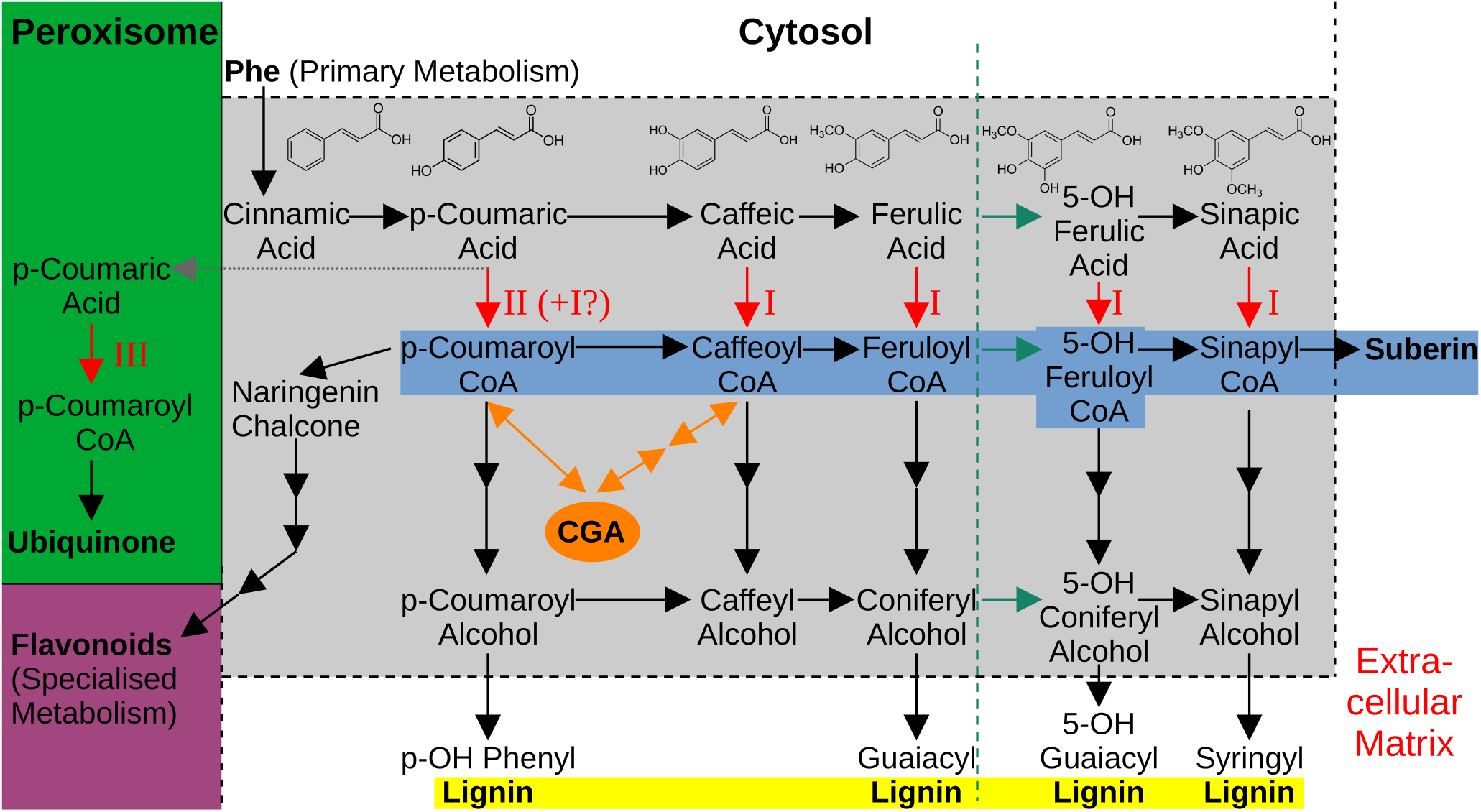
Model of the Phenylpropanoid Metabolism as Intermediate Metabolism with Departing Fluxes and CGA pool. The phenolpropanoid metabolism is inside the grey box. Arrows represent reactions: the grey dotted arrow reflects transport over the peroxisomal membrane; red arrows reflect 4CL catalysed reactions; green arrows reflect ferulate 5-hydroxylase catalysed reactions restricted to angiosperms. The green line indicates as such a barrier in gymnosperms. Indicated are the canonical 4CL classes I and II as well as the suggested peroxisomal class III. Four major fluxes are distinguished: ubiquinone (green); flavonoid (purple); lignin (yellow) and suberin (blue), the latter two directing at extracellular matrix polymers. CGA is considered as the major phenolic antioxidant pool, hence the double arrows.

Phenylalanine is the common precursor of the phenylpropanoid metabolism; it is converted to subsequently cinnamate and p-coumarate by phenylalanine ammonia lyase and cinnamate 4-hydroxylase. p-Coumarate is then either converted to caffeate or 4-coumaroyl-CoA, or it is transported to the peroxisome where a peroxisomal 4CL converts it to 4-coumaroyl-CoA in order to feed ubiquinone biosynthesis (Block et al. 2014). Cytosolic 4-coumaroyl-CoA feeds the flavonoid pathways and the formation of CGA. Cytosolic CL enzymes also catalyse the activation of several other hydroxycinnamic acids (Figure 1) into their corresponding CoA esters (Knobloch and Hahlbrock 1975, 1977). These activated phenolic acids are precursors for the poly-aromatic domains of suberin and lignin, the latter via monolignols. All together, we consider four major fluxes towards ubiquinone; suberin; lignin; and flavonoids, all fed by phenylalanine or the CGA pool (Figure 1).

Given its involvement in ubiquinone and suberin production, the evolution of the phenylpropanoid pathway, and more specifically the 4CL superfamily, marks the origin of land plants. Gymnosperms and angiosperms appear to have evolved somewhat differently, resulting in differences in lignin and suberin composition. Gymnosperms lack ferulate 5-hydroxylase (Figure 1) and cannot produce syringil precursors (Zhao et al. 2010), their suberin appears to contain mostly ferulate and no detectable sinapate (Barros et al. 2023). They also rely more on terpenes and resin as rapid wound response. Flavonoids are absent in early-diverging lineages such as algae, pteridophytes, lycophytes and bryophytes and developed later, having more specialised roles including protection against ultraviolet light and signalling in plant-microbe interactions (Koes et al. 1994). Both angiosperms and gymnosperms produce a wide range of flavonoids, including the colourless anthocyanidins, but gymnosperms do not produce the colourful anthocyanins, which are important in the pigmentation of flowers, fruits and seeds to attract pollinators and seed dispersers. The presence of anthocyanins appears to correlate with the greater diversification of angiosperms. Hence, it appears the variation in hydroxycinnamates is an important aspect of the phenylpropanoid metabolism and its downstream fluxes.

4CL enzymes are encoded by an intermediately complex gene family: Arabidopsis and rice contain over 10 different isoforms annotated as either 4CL or 4CL-like in the SwissProt database. The 4CL family is part of the Acyl-CoA Synthetase (ACS) family, which on its turn is part of the ANL superfamily of adenylating enzymes (Babbitt et al. 1992). This derives its name from its three constituent families: the ACS family; the Non-ribosomal peptide synthetases; and the Luciferase family (Gulick 2009), which share the adenylation reaction as part of otherwise diverse reactions. The presence of a putative AMP-binding domain with a conserved motif named Box I [LIVMFY]-X-X-[STG]-[STAG]-G-[ST]-[STEI]-[SG]-X-[PASLIVM]-[KR]) is used as an important computational criterion for establishing whether a protein belongs to the ANL superfamily (Fulda et al. 1994). Superfamilies are hierarchic, which makes classification both important and difficult. As such, others have designated Box II, or the GEICIRG motif, as signature for 4CL enzymes (Becker-André et al. 1991). Indeed, only the Arabidopsis homologues annotated as 4CL, and not 4CL-like, have the GEICIRG motif (Costa et al. 2005).

The determination of crystal structures of 4CL enzymes from *Populus tomentosa* (Hu et al. 2010) and tobacco (Li and Nair 2015) in two conformations enabled the dissection of 4CL catalysis. The initial adenylation, performed in the adenylation conformation and the presence of ATP and MgZ⁺, results in a transient hydroxycinnamate-AMP anhydride. A significant movement of the C-terminal part of the enzyme results in the thioester-forming conformation that allows binding of the second substrate CoA, followed by a nucleophilic attack on the carbonyl carbon of the acyl-adenylate by the CoA phosphopantetheine-thiol.

Cytosolic 4CL enzymes are divided into Class I and Class II (Ehlting et al. 1999). Class I homologues are involved in the biosynthesis of suberin and lignin, as their expression is induced by wounding and confined to stems and roots (Ehlting et al. 1999; Cukovic et al. 2001),whereas Class II homologues are associated with flavonoid biosynthesis, as they are mainly expressed in pigmented tissues and induced by UV treatment. Correspondingly, different substrate preferences for Class I and Class II enzymes have been hypothesised. Differences in substrate preference among Class II and various Class I isoforms have been reported *in vitro* for *A. thaliana* (Costa et al. 2005), rice (Gui et al. 2011), and soybean (Lindermayr et al. 2003), but offered no clear relationship between functional class and substrate preference. Class I At4CL4 is the only isoform that prefers ligating sinapate, the most voluminous hydroxycinnamate. This was related to a deletion and susbstantiated by mutant analysis in soybean (Lindermayr et al. 2003). Interestingly, Brassicaceae appear to produce family-specific sinapate esters (Milkowski et al.).

Only two nearly identical Class I isoforms have been described in potato (Becker-André et al. 1991). Hence, in order to study 4CL in potato tubers, the repertoire of 4CL must be properly analysed. BLAST (Altschul et al. 1990), typically used to allow transitive function assignation, can only assign a general function to a protein and cannot reliably identify whether a given homologue is a Class I or Class II enzyme. Furthermore, many homologues come with the non-specific annotation of 4CL-like. For instance, an initial *in silico* analysis revealed that the *Arabidopsis* genome has 14 genes annotated as putative 4-coumarate-CoA ligases. Eleven of these were expressed heterologously and tested for activity. Only four were catalytically active *in vitro* towards known 4CL hydroxycinnamate substrates, confirming that the 4CL protein family in *Arabidopsis* has at least four members. The remaining seven enzymes were designated as 4CL-like proteins (Costa et al. 2005). Importantly, this finding was in agreement with previous phylogenetic analyses (Shockey et al. 2003; Costa et al. 2003). The enzymes encoded by several 4CL-like genes were mostly shown to be ACS that have medium- to long-chain fatty acids or more complex carboxylates as substrate (Schneider et al. 2005; Kienow et al. 2008). In particular, it was demonstrated that the *Arabidopsis* 4CL-like gene At1g62940 is a medium- to long-chain fatty ACS required for sporopollenin monomer biosynthesis in tapetal cells, designated ACOS5 (De Azevedo Souza et al. 2009). 4CL-like genes At5g63380, At4g05160, At1g20500 and At1g20510 encode peroxisomal enzymes that have the capacity to activate some Jasmonic Acid (JA) precursors such as cyclopentenone 12-oxo-phytodienoic acid *in vitro* (Schneider et al. 2005). Particularly, At1g20510 has been designated as OPC-8:0-CoA ligase (OPCL), using 8-[(1R,2R)-3-oxo-2-[(Z)-pent-2-en-1-yl]cyclopentyl]octanoic acid (OPC-8:0) as substrate (Koo et al. 2006). More recently, 4CL activity with kinetics similar to cytosolic 4CLs was reported for a peroxisomal 4CL-like protein (At4g19010) involved in ubiquinone biosynthesis (Block et al. 2014), a finding important for the definition of the 4CL/ACS superfamily we intend to investigate. Another striking aspect we identified is that *A. thaliana*, rice and soybean have more than a single Class I homologue. Since there are different substrates for Class I enzymes (Figure 1), we suspected that Class I enzymes may have additional subclasses, despite the fact that no clear substrate preference has been demonstrated *in vitro*.

The original aim of the present work was to study the 4CL enzymes in *Solanum tuberosum* tubers in order to define which gene or genes are involved in flavonoid biosynthesis and which in lignin and suberin production. Given the complex background of the 4CL/ACS superfamily as well as some spurious data in literature and outdated SwissProt sequence annotations, we performed a thorough classification of the 4CL/ACS superfamily with the objective to identify and describe the functionally different subfamilies, as of here referred to as functional (sub)families. In section 1 we define and cluster the 4CL/ACS superfamily using a thoroughly mined, representative sequence set. In section 2, we study the differences between the major subfamilies. In section 3, we used the obtained partition of the representative sequence set as a training set to classify a larger sequence set from 200 complete proteomes to show how the cytosolic 4CL enzymes evolved, with a focus on the classification and functional diversification of Class I enzymes. Finally, in section 4 we analyse public RNA-seq data in potato and the expression of Class I and II 4CL encoding genes upon wounding and in coloured and white potato tubers.

## Materials and methods

### Plant material

*S. tuberosum* ssp. *andigena* varieties (Table 1) were grown in fields located in Quebrada de Humahuaca, Jujuy, Argentina during the 2010/2011, 2011/2012 and 2016/2017 campaigns. All varieties were planted on the same date in random plots and harvested at the end of their respective cycles. We used a single batch from the 2010/2011 campaign, and two independent batches from the 2011/2012 campaign for each variety. Skin and flesh from ten freshly harvested tubers were pooled to generate a representative sample. The material was immediately frozen in liquid nitrogen, lyophilized and stored at -80 °C until analysis.

**Table 1.** Description of the potato varieties used in this study. Indicated are colours according to the following codes: Y, yellow; C, cream; PK, pink; R, red; P, purple. Lowercase letter refer to a secondary colour.

| Variety | Colour |  | Relative Phenolic Content |  |
| --- | --- | --- | --- | --- |
|  | Flesh | Skin | Flesh | Skin |
| Chaqueña | Cpk | PK | Intermediate | Intermediate |
| Santa María | R | R | High | High |
| Waicha | Y | PK | Low | Intermediate |
| Moradita | Y | P | Low | High |

### Wound experiment

Tubers from two independent batches of *S. tuberosum* spp. *andigena* cv. Santa María (2016/2017 campaign) were peeled and cut into 2-3 mm thick slices. Peels represent native periderm and were frozen in liquid nitrogen, lyophilized and stored at -80°C. Slices were left to heal at room temperature in saturated humidity and darkness for 0 and 72h and then frozen in liquid nitrogen, lyophilized and stored at -80 °C. Anthocyanin content was measured by the pH differential method (Durst and Wrolstad 2001).

### RNA extraction, cDNA synthesis, primer design, and qRT-PCR

RNA was extracted from tuber samples using the cetyltrimethylammonium bromide method (Li et al. 2011). cDNA was synthesized using 2 µg of total RNA, pretreated with RNase-free DNase I (Invitrogen, Carlsbad, CA), anchored oligo(dT) 15 VN primers and MMuLV reverse transcriptase according to the manufacturer’s description (Invitrogen).

Isoform-specific primers (Supplemental Table 1) were designed to anneal in the 3’ coding region of St4CL genes flanking the last variable-length intron (Supplemental Figure 1A) such that amplicon length discriminates RNA from gDNA origin. The amplicon length of PCR products using as a template gDNA from *Solanum phureja* confirmed primer specificity (Supplemental Figure 1B).

Relative transcript levels were determined by qRT-PCR in a 10 µL reaction volume with 20 ng RNA equivalent cDNA, 300 nM gene-specific primers and 5 µL SYBR Green Mix (Roche, Mannheim, Germany). Amplification was done using StepOneTM Real-Time PCR System according to the manufacturer’s description (Applied Biosystems, Foster City, CA). Relative expression levels were calculated by the ΔCT method (Livak and Schmittgen 2001) by normalizing the CT levels of target genes to the geometric mean of CT levels of the housekeeping gene encoding cytoplasmic ribosomal protein L2.

### RNA-seq analyses

Transcript analysis of 4CL/ACS potato gene family from publicly available RNA-seq data was performed using the Potato eFP Browser (Winter et al. 2007) using standard settings and the datasets provided by the PGSC (Massa et al. 2011). Note that these do not include data for StUACSII.1, StUACSIII, and StACOS5 since these were not included in PGSC v4.03.

### Sequence identification and analysis

All multiple sequence alignments (MSA) were performed using MAFFT (Katoh and Standley 2013; Katoh et al. 2018) G-INS-i at the MAFFT website (Katoh et al. 2019) or stand alone using the command *mafft --globalpair --maxiterate 16 <sequences.fsa> > <sequences.faa>*.

Phylogeny was performed on BMGE-trimmed (Criscuolo and Gribaldo 2010) MSAs (*java -jar BMGE.jar -i sequences.faa -t AA -h 0.8 -of sequences08.faa*) using PhyML (Guindon et al. 2010) with default settings (automated model selection resulting in custom model and four rate classes). We used FastTree (Price et al. 2010) with 1000 bootstraps using Booster for statistical support (Lemoine et al. 2018). Graphical representations of phylogenies were made using Dendroscope (Huson and Scornavacca 2012).

Sequence mining concerned three datasets. First, an initial training-set of sequences was obtained by a specific PHI-BLAST (Zhang et al. 1998) using the PROSITE (Sigrist et al. 2010) PS00455 consensus sequence as pattern and At4CL1 as well as Os4CL1 as queries against the UniProtKB/SwissProt database restricted to plants (taxid:3193) with word-size 3. Sequences with an Expect-value ≤ 1e-50 were selected and manually inspected for partial sequences. The accepted sequences were used to perform an iterative HMMER (Eddy 2011) based sequence mining using a compiled fasta file containing all the complete proteomes of 15 selected species (Table 2). In short: an MSA is translated by *hmmbuild* to a hidden Markov model. The compiled fasta file was screened by *hmmsearch,* using a specific cut-off threshold determined by the lowest scoring sequence of the input set. Positive sequences were added to the initial training set providing a novel set for *hmmsearch*. The obtained sequences were subjected to a default Seqrutinator (Amalfitano et al. 2024) filter removing partial sequences and sequences that otherwise likely do not encode a functional homologue. Sequence, XP_006366955 from *S. tuberosum* was obtained from the NCBI database. Redundant sequences showing 100% identity were eliminated using CD-HIT-2D (Li and Godzik 2006)

**Table 2.**
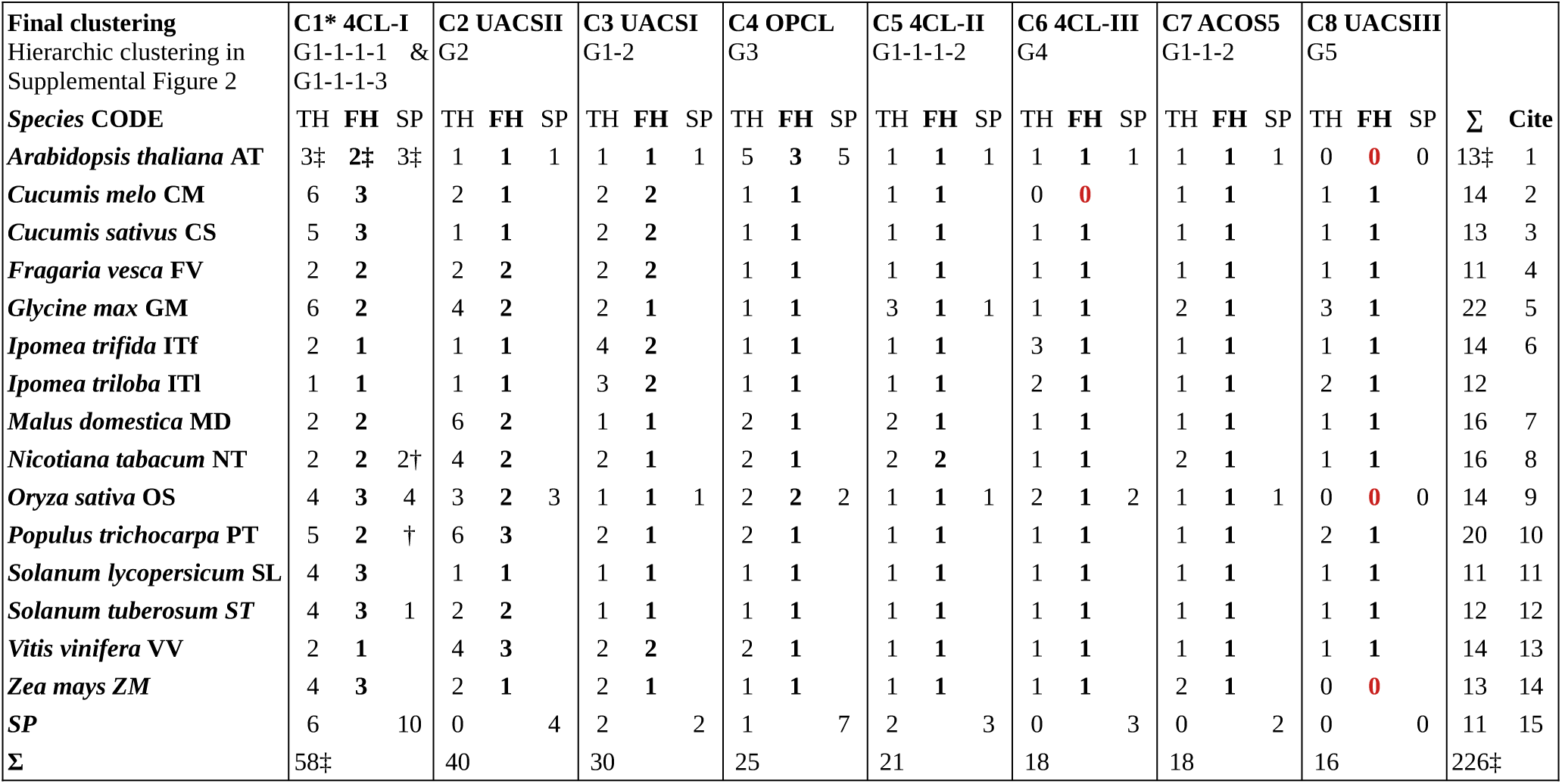
Total Number of Homologues and Functionally Non-redundant Homologues per Species and Cluster. Sequences were obtained from complete proteomes from 15 indicated species, SwissProt, or PDB. TH: Total number of Homologues, FH: Functional (Non-Redundant) Homologues; SP: SwissProt: The SP row shows the number of entries from other plant species (TH column) and the number of entries from the 15 selected species. Missing homologues are in red. *: C1 consists of subclusters C1a and C1b as shown in Figure 2. ‡: This number includes sinapate:CoA ligase sequence Q9LU36, absent in the tree of Figure 2. † Sequence with X-ray structure (PDB). UACS: Uncharacterised ACS; OPCL: OPC-8:0 CoA ligase; ACOS5; Acyl-CoA Synthetase; 4CL-I, 4CLII and 4CL-III: Class I, II and III (peroxisomal) 4CL. Cites: 1 (Costa et al. 2003); 2 (Garcia-Mas et al. 2012); 3 (Huang et al. 2009); 4 (Shulaev et al. 2011); 5 (Schmutz et al. 2010); 6 (Wu et al. 2018); 7 (Velasco et al. 2010); 8 (Edwards et al. 2017); 9 (Kawahara et al. 2013); 10 (Tuskan et al. 2006); 11 (Lin et al. 2014); 12 (Xu et al. 2011); 13 (Jaillon et al. 2007); 14 (Schnable et al. 2009); 15 (Bairoch and Apweiler 2000).

Shannon sequence and sequence-like logos were made with the representative sequence set of 225 sequences and in house scripts, applying logomaker (Tareen and Kinney 2020). Structures were obtained from the PDB database, models were obtained from UniProt (Consortium 2017). Images of proteins structures were made using PyMOL (Schrödinger, LLC 2015), composite images were made using GIMP.

## Results

### 1 The Phylogeny of the 4CL/ACS Superfamily reveals Eight Functional Subfamilies

We selected 42 sequences (*Repo-Blast4CL.fsa*) from a PHI-BLAST search (Zhang et al. 1998) against the SwissProt database (Bairoch and Apweiler 2000) and aligned them into an MSA (*Repo-Blast4CL.faa*) that we used to construct a HMMER profile (Eddy 2011) (*Repo-Blast4CL.hmm*). With this profile we initiated a dynamic screening of complete plant proteomes of the 15 plant species listed in Table 2, and Protein Data Bank (PDB) (Berman et al. 2000) for homologues with structure information. Notably, all homologues from *A. thaliana* and *O. sativa* are represented in SwissProt, mostly without experimental validation. After two iterations we reached sequence convergence. Manual curation removed partial, erroneous, or redundant entries, yielding a preliminary set of 226 sequences, which we used to generate an MSA and reconstruct a maximum likelihood phylogenetic tree.

We subjected the tree and corresponding sequences to HMMERCTTER’s (Pagnuco et al. 2018) similarity-based clustering to identify potential functional or taxonomic groups. HMMERCTTER identifies monophyletic clades as clusters, and constructs cluster-specific MSAs and corresponding HMMER profiles. Then, it determines if the clusters are 100% precision and recall in a hmmsearch self-detection (100% P&R-SD) by testing if all cluster sequences score higher than all non-cluster sequences. This approach potentially separates functional families: monophyletic (sub)families with different functional aspects, since similar functional constraints acting on paralogues tend to conserve the sequences, yielding higher *hmmsearch* scores.

Initial clustering of the preliminary dataset identified *A. thaliana* sequence with SwissProt identifier Q9LU36 as a sequence outlier that severely disturbed automated clustering. The sequence corresponds to the 4CL that most efficiently converts sinapate *in vitro* (Costa et al. 2005), This different substrate specificity suggests a rare trajectory towards neofunctionalization relative to the class I 4CLs. Therefore we temporarily separated this sequence; in section 3 we will show this sequence indeed represents a novel, Brassicales-specific subfamily. The novel sequence set (*Repo-4CLACS_Representative.fsa*) was aligned (*Repo-4CLACS_Representative.faa*), used for phylogenetic reconstruction (*Repo-4CLACS_Representative.nexml*) and could be successfully clustered by HMMERCTTER into nine clusters (Figure 2).

**Fig 2.**
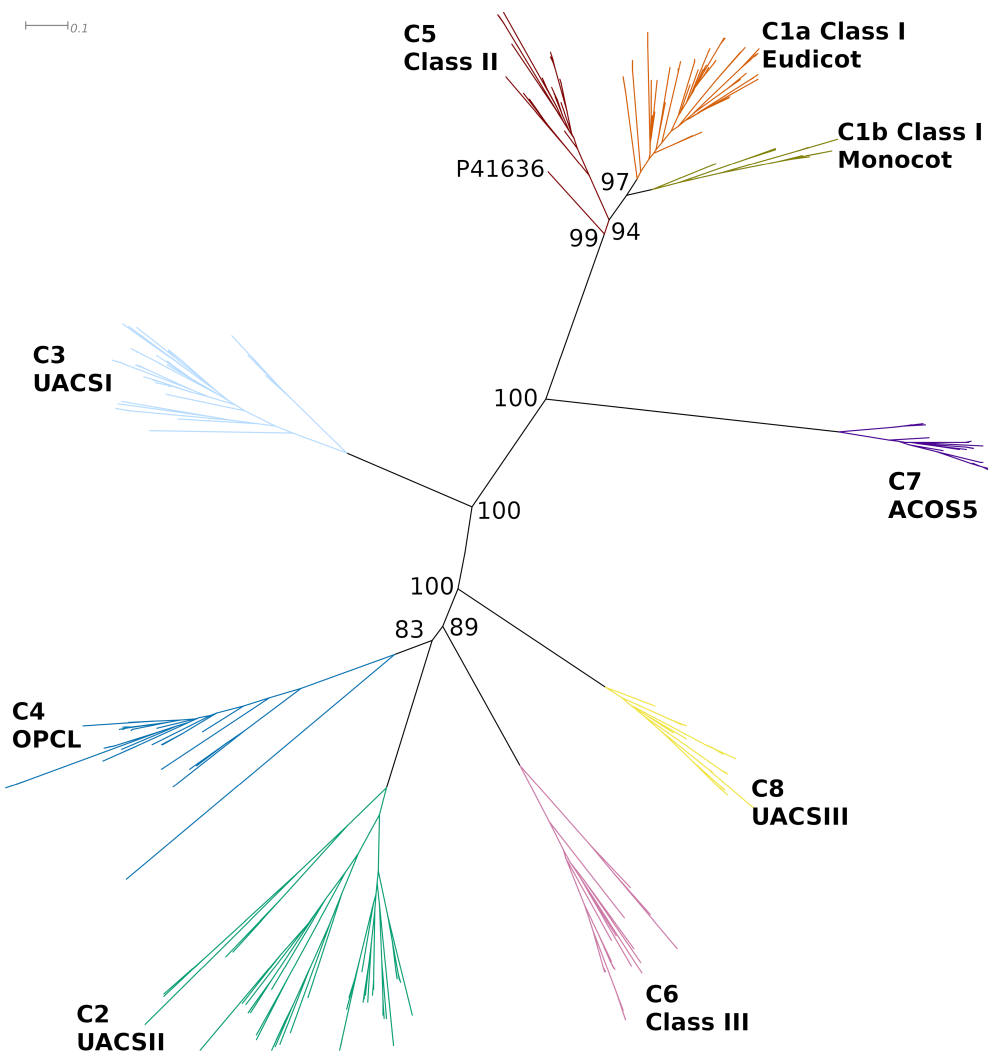
Maximum Likelihood Tree and Final Partition of Representative 4CL/ACS Sequences. Clades with different colours are 100% P&R-SD. C1a and 1b, representing Class I sequences are not 100% P&R-SD as single functional cluster but only as taxonomically split clusters. Numbers are according to cluster size, taking C1 as the combined 1a and b. The codes indicate consensus classification (Class I and II), designated functional classification (Class III, OPCL and ACOS5), or temporary designation as Unclassified ACS (UACSI to III). P41636 is a 4CL homologue from gymnosperm P. taeda*, and makes the CII class paraphyletic in this tree. Numbers indicate bootstrap support of major bifurcations. The scale bar shows the number of amino acid substitutions per site reflecting evolutionary distances of the different taxa. A bootstrapped tree in Newick format is in Supplemental Document 1*.

The initial clustering of the representative dataset contained, besides four compact, peroxisomal sequence clusters, a large cluster with a peroxisomal subcluster; a single well separated cytosolic subcluster; and a complex, cytosolic cluster with both Class I and Class II 4CL sequences at sequence distances that are large as compared to the distance between ancestral and basal nodes. The four compact clades have single *functional non-redundant homologues (FH)* for most species (Table 2). The FH metric is defined as a set of one or more paralogues that emerged very recently. They occur as near identical sister sequences in the phylogenetic tree and represent functionally redundant rather than diversified homologues. As such, the FH metric forms a good metric to identify functional subfamilies in a hierarchic clustering. The designation of functional family is further substantiated by relatively large distances between the clades’ ancestral and basal nodes.

The large cluster required hierarchic subclustering. In terms of evolution, each subsequent subclustering corresponds to an *inception*; a commencement of a novel layer of functionality. A total of three subsequent subclustering of such inceptions, as shown in Supplemental Figure 2, was required to pinpoint the Class I functional family. Separation of Class I from Class II homologues automatically resulted in Class I subclusters for eudicot and monocot homologues. This has resulted from evolution that was guided by a combination of functional constraint and supposedly more neutral taxonomic drift. This combination of natural selection and drift occurred in all lineages, often but not always resulting in taxon specific clades, and forms a well known and important issue in protein superfamily analyses.

Most clusters shown in Figure 2 appear well separated from their basal node and show a single FH for most species. Exceptions include cluster C8, which appears specific to dicotyledons and lacks an Arabidopsis homologue, likely reflecting a gene birth event in the eudicot lineage and a subsequent loss in the *Brassicales* ancestor, according to Nei’s birth and death model of evolution (Nei et al. 1997). Conversely, *A. thaliana* possesses five relatively divergent paralogues in C4. *Cucurbita melo* appears to lack a C6 homologue, likely reflecting an incomplete genome sequence rather than a gene loss. The major discrepancy is formed by the Class I cluster showing two or more FH for most species. Also the UACSII clade has relatively many species with two or more FH.

The goal of this functional classification is to allow accurate transitive sequence function assignation. Besides assigning the well described Class I and Class II enzymes, we designated C6 homologues as Class III sequences since the *A. thaliana* orthologue was unequivocally shown to be the major peroxisomal 4CL required for ubiquinone biosynthesis (Block et al. 2014), forming a third class of 4CL activity. Similarly, C7 represents the ACS designated ACOS5 based on the well described activity in *A. thaliana* (De Azevedo Souza et al. 2009). Note that both sequences are still annotated as “4CL-like” in the SwissProt database. Then, many *A. thaliana* sequences in C4 have been related to OPCL activity, either by means of *in vitro* activity (Kienow et al. 2008) or by means of T-DNA mutants (Koo et al. 2006). Thus we designated the C4 clade as OPCL. For the remaining clades we found no clear evidence in literature to support functional assignation. These subfamilies are clearly under a functional constraint, hence do form different functional classes and were designated as Uncharacterised ACS (UACS I to III) (Table 2). The corresponding HMMER profiles, cluster specific ID files and the numerical output of a 100% P&R-SD screen for all subfamilies are available for sensitive and precise HMMERCTTER or manual 4CL/ACS sequence mining in the data repository.

### 2 Sequence Analysis Shows Few Subfamilies Display Clear Differences in Their Binding Pockets

We used the described sequence hallmarks of Box I and Box II to examine potential differences among the various subfamilies. Figure 3 presents sequence logos for both Boxes across all eight functional clusters, with taxonomic clusters 1a and 1b, representing Class I cytosolic 4CL, combined. Both motifs appear highly conserved, confirming their utility as hallmarks for the 4CL/ACS superfamily. Our analysis revealed several substitutions within or near the pyrophosphate-binding or P loop, part of Box I and described in detail by Li and Nair (Li and Nair 2015). This loop is more or less enclosed by P187 and P196 that are substituted by L and S or V, respectively (Figure 3a&b). Since proline induces structural turns, positions 187 and 196 may thus shape or affect the dynamics of the P loop, depending on the subfamily specific amino acid. To assess the structural implications of these substitutions, we performed comparative structure analysis using structures from PDB and Alphafold2 models provided by UniProt (Figure 3c). The most pronounced deviation was observed in the C2 model, whose P loop adopts a conformation similar to that of the thioesterification form of 5BST. P187 appears to break up a predcited β-sheet in C2 (Figure 3c). S189 appears in contact with the co-crystallised substrate of 5BST.

**Fig 3.**
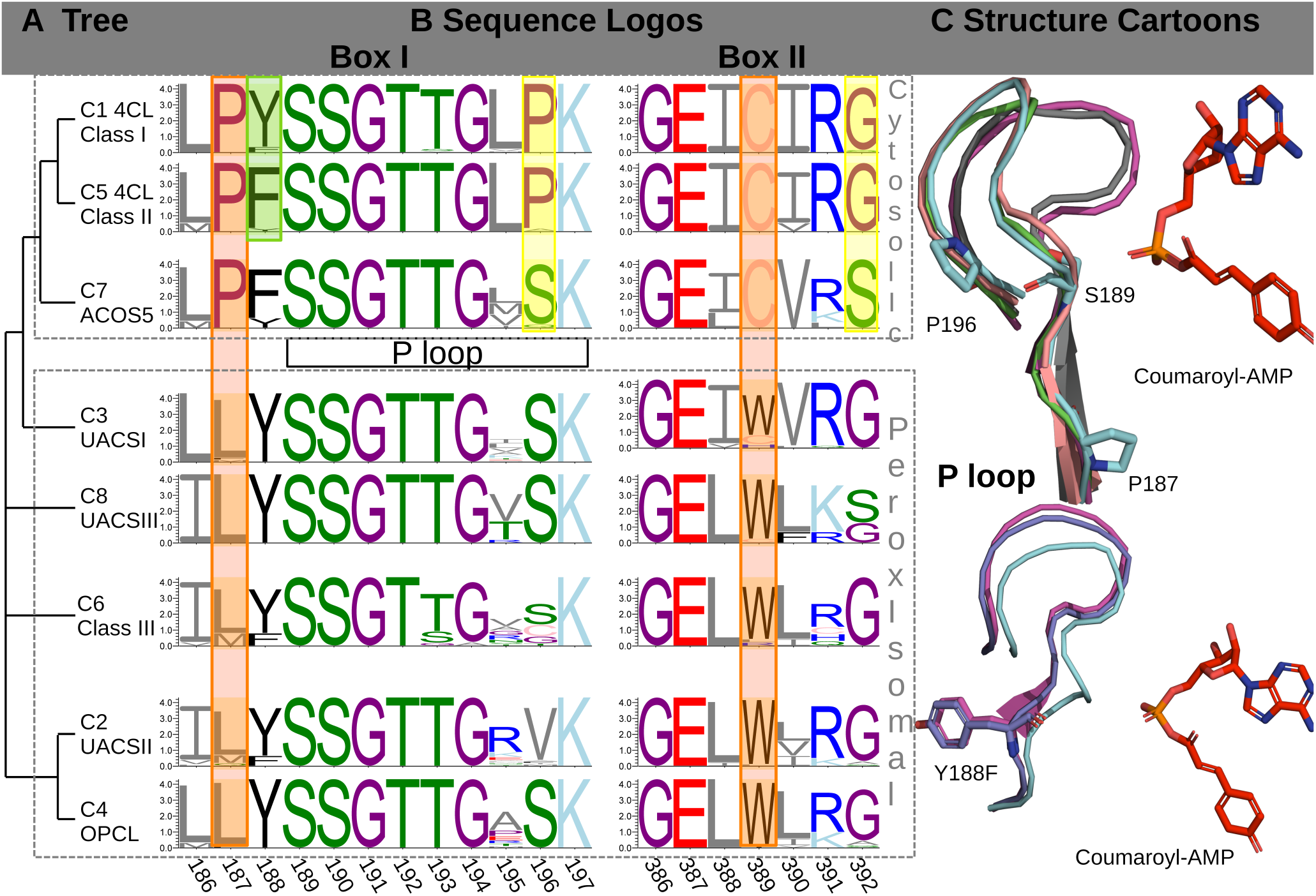
Cluster Dependent Differences in Box Motifs. **A Tree Topology with Clustering Aligned with Sequence Logos in B.** Box I contains the P loop. Numbers according to 5BSM, which is the only structure iin adenylation-conformation. Logo placement according to the most likely hierarchic evolutionary pathway, as indicated in A. Colours: Green: Polar STQN; Blue: Basic RH; Cyan; Amphipatic K; Grey: Small Hydrophobic LIVMA; Black: Large Hydrophobic FYW; Red: Acid DE;, Purple: Structure PG, Pink: Redox C. Highlights point out differences: yellow 4CL vs. ACOS5; green Class I vs. Class II; orange between cytosolic and peroxisomal subfamilies. **C Cartoon Showing P Loop Displacement (Top) and Y188F Substitution (Bottom).** 5BSM is in cyan with P187 and P196 shown in sticks (Top). 5BST is in cyclamen with substrate analogue coumarate-AMP and S189 in sticks (Top); Pink: C3 (P187L-P196S, Q9M0X9); Green: C7 (P187-P196S, Q7XXL2); Grey: C2 (P187L-P196V, Q84P23). Y188 (5BSM) and F188 (C5, purple A0A2H5AIY4) are shown in sticks (Bottom).

Substitution Y188F distinguishes Class I from Class II enzymes. The Class II model (A0A2H5AIY4) places its phenylalanine at the same location as Y188 in a thioesterification conformer (Figure 3c). Because this residue points away from the substrate, there is no direct electrostatic interaction with the substrate, suggesting that this substitution contributes to fine-tuning P-loop dynamics rather than to substrate affinity.

Box II corresponds to the A6 motif and has been proposed to play primarily a structural role, located relatively close to the substrate (Gulick 2009). It shows two major substitutions among the subfamilies (Figure 3b). Cytosolic sequences carry C389 where peroxisomal sequences have W389, forming a rare conserved substitution that, according to Alphafold models provided by UniProt, has surprisingly little structural impact. Similarly, the more subtle G392S substitution, found between cytosolic and most peroxisomal sequences, appears not to affect overall conformation.

While the descriptions of Box I and II are valuable for a global sequence-based characterization, structural motifs are likely more informative in explaining substrate specificity. In particular, the relatively small sequence divergence between the 4CL and ACOS5 subfamilies shown in Box I and Box II, seems insufficient to explain the large difference between coumarate and oleate, alleged substrates of 4CL and ACOS5 respectively. We therefore used the Nt4CL2 structures to define approximate binding pockets. Note that 5BSM, co-crystallized with ATP, represents the adenylate-forming conformation, whereas 5BSR, 5BST, 5BSU, and 5BSV represent the thioesterification conformation. All residues within 5 Å of the co-crystallized substrates described by Li and Nair (Li and Nair 2015) were identified as part of the binding pocket (Supplemental Table 2). Sequence-like logos were then generated separately, for residues forming the hydroxycinnamoyl-AMP/ATP pocket (Figure 4a) and for those contacting the CoA moiety (Figure 4b).

**Fig 4.**
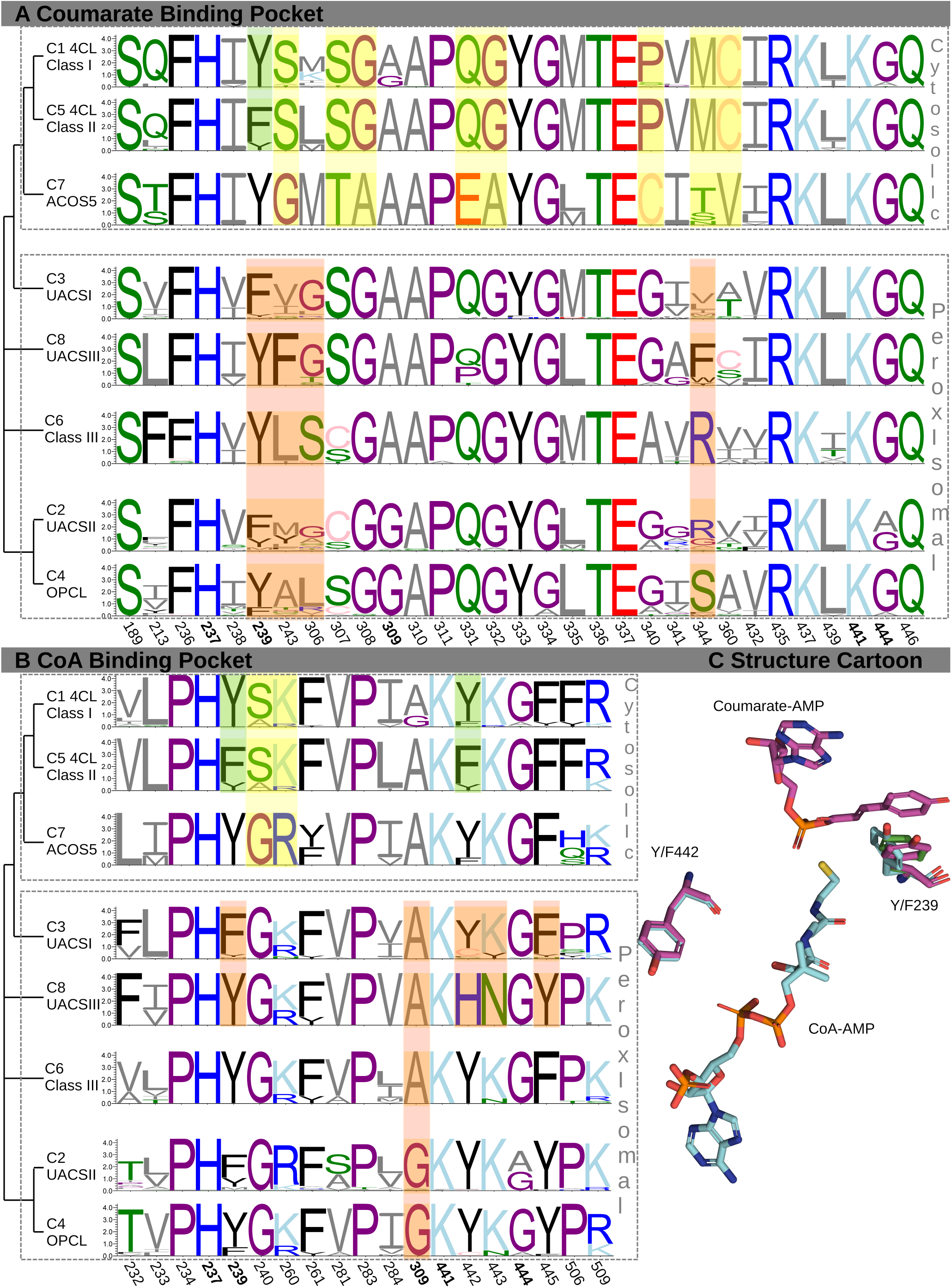
Cluster dependent differences in binding pockets. **A Logos of coumarate binding pocket in eight clusters. B Logos of the CoA binding pocket in eight clusters. C Cartoon of the Y239F and Y442F substitutions near the ligands.** Numbers according to 5BSM, underlined position takes part in Box I, boldface positions are in both pockets. Logo placement according to the most likely hierarchic evolutionary pathway, as indicated on the left. Colours: Green: Polar STQN; Blue: Basic RH; Cyan; Amphipatic K; Grey: Small Hydrophobic LIVMA; Black: Large Hydrophobic FYW; Red: Acid DE;, Purple: Structure PG, Pink: Redox C. Highlights point out differences: yellow 4CL vs. ACS: green Class I vs. Class II; orange among different peroxisomal subfamilies.

An interesting observation is that 4CL and ACOS5 enzymes differ by eight substitutions among the 31 residues (*∼*25%) forming the hydroxycinnamoyl pocket, confirming that major substrate differences require more extensive changes than the small P-loop displacement seen in Figure 3c. Merely two conserved differences were identified in the 19-residue CoA-binding pocket (*∼*10%). The ACOS5 binding sites do not only differ from the binding sites of cytosolic 4Cl enzymes, they are the most distant overall. In fact, the binding sites of the peroxisomal enzymes are surprisingly similar to that of cytosolic 4CL enzymes. One the one hand it fits with Class III 4CL activity, on the other it does not easily explain OPCL activity on OPC-8:0, a rather different substrate. The only clear difference is that OPCL and the related UACSII have the A309G substitution, part of both binding pockets (Figure 4), which may accommodate more easily the somewhat more voluminous JA precursors.

In contrast, few substitutions were observed between Class I and Class II enzymes, consistent with more closely related substrate profiles. One difference, Y239F, lies at the interface of both pockets. This residue preserves a six-carbon aromatic ring that aligns with the pyranose of the coumarate moiety (Figure 4c) suggesting a π–π interaction, as previously proposed by Li and Nair (Li and Nair 2015). Indeed, Y239F and Y239A mutants of Class I Nt4CL2 displayed approximately five- and 150-fold increases, respectively, in the K_m_ for coumarate. Interestingly, since this position appears to influence both pockets, it may also stabilize the transition state.

Another substitution is Y442F, which only takes part in the CoA binding pocket (Figure 4c). The residues appear to align perfectly but the absence or presence of the tyrosine-hydroxyl may contribute to CoA binding. A further notable set of substitutions at the same position is Y442H-K443N, observed in C8. K443 in 5BSR forms one of few hydrogen bonds with CoA, and its substitution to alanine was shown to have only minor kinetic effects (Li and Nair 2015). Together, these adjacent substitutions highlight evolutionarily flexible yet functionally critical positions within the superfamily, likely evolving in concert with metabolic context. Computational analyses focused on identifying specificity-determining positions may reveal additional diversifying features among peroxisomal subfamilies, which goes beyond the scope of this work.

### 3 Dedicated Phylogeny of the 4CL Family Reveals a Diversification of Class I into a Complex Set of Subfamilies

Two aspects described in section 1 require further investigation. First, most species have more than a single Class I FH, which allows for functional redundancy and diversification. We hypothesise Class I constitutes more than a single functional family, exemplified by the sinapate:CoA ligase from *A. thaliana.* Visual analysis of the Class I clade in Figure 2 does, above all, reveal divergence that occurred as a result of taxonomic drift between eudicot and monocot sequences clustering separately in clades 1a and 1b. We mined complete proteomes of 200 additional plants for the presence of 4CL/ACS homologues and reconstructed a cytosolic 4CL enzyme phylogeny with higher resolution, in search for additional clustering patterns. This allowed to look into the disputed position of SwissProt entry P41636 from gymnosperm *P. taeda* (Figure 2) by including two gymnosperms, a fern, a bryophyte and a lycophyte as non-angiosperm outgroups. Inclusion of such outlier sequences may help clarify the nature of the *P. taeda* sequence and may help pinpoint the ancestral 4CL enzyme showing when the Class I - Class II diversification occurred.

We obtained sequences of all putative homologues using the general HMMER profile described in section 1, and complete proteomes of 200 species listed in Supplemental Document 2. This included the removed sinapate:CoA ligase sequence from *A. thaliana,* alongside several closely related Brassicales homologues. Seqrutinator (Amalfitano et al. 2024) removed sequences that are partial or otherwise likely not functional. Sequences accepted as functional homologue were then classified using the nine cluster-specific HMMER profiles obtained with the training set (*Repo-C*.hmm*) and HMMERCTTTER’s classification module (Pagnuco et al. 2018).

We performed a series of control analyses to show the performance of the classification procedure. For instance, all hitherto designated functional families showed mostly a single FH for most species (Supplemental Document 2). Specifically, UACSIII, the clade which we had previously reported as lacking monocot and Arabidopsis homologues, actually lacks Poales and Brassicaceae homologues. We also found that there are relatively few peroxisomal gymnosperm sequences, restricted to clades C2 and C4, whereas the only and single peroxisomal homologues for the fern *Azolla filiculoides* and lycophyte *Selaginella moellendorfii* were identified in C3. No sequences for bryophyte *Physcomitrella patens* were found in any peroxisomal clade nor in the C7/ACOS5 clade. This last clade appears to have a single candidate for all other species we included (Supplemental Document 2).

Next we focused on the sequences identified by the three profiles representing Class I sequences (*Repo-C1a.hmm* and *Repo-1b.hmm*) and Class II sequences (*Repo-C5.hmm*). The sequences were combined (*Repo-4CLSequences.fsa*), aligned and used to reconstruct a phylogenetic tree representing the evolution of the cytosolic 4CL family in plants. The obtained tree (Figure 5a; *Repo-4CLTree.nexml*) shows clustering with taxonomic grouping, suggesting further functionalisation. Unfortunately, HMMERCTTER could not cluster this sequence set properly, given the combination of short inter-clade and large intra-clade distances and the presence of the SL homologues. We therefore applied intuitive clustering and analysed the taxonomic distribution in order to identify clustering patterns.

**Fig 5.**
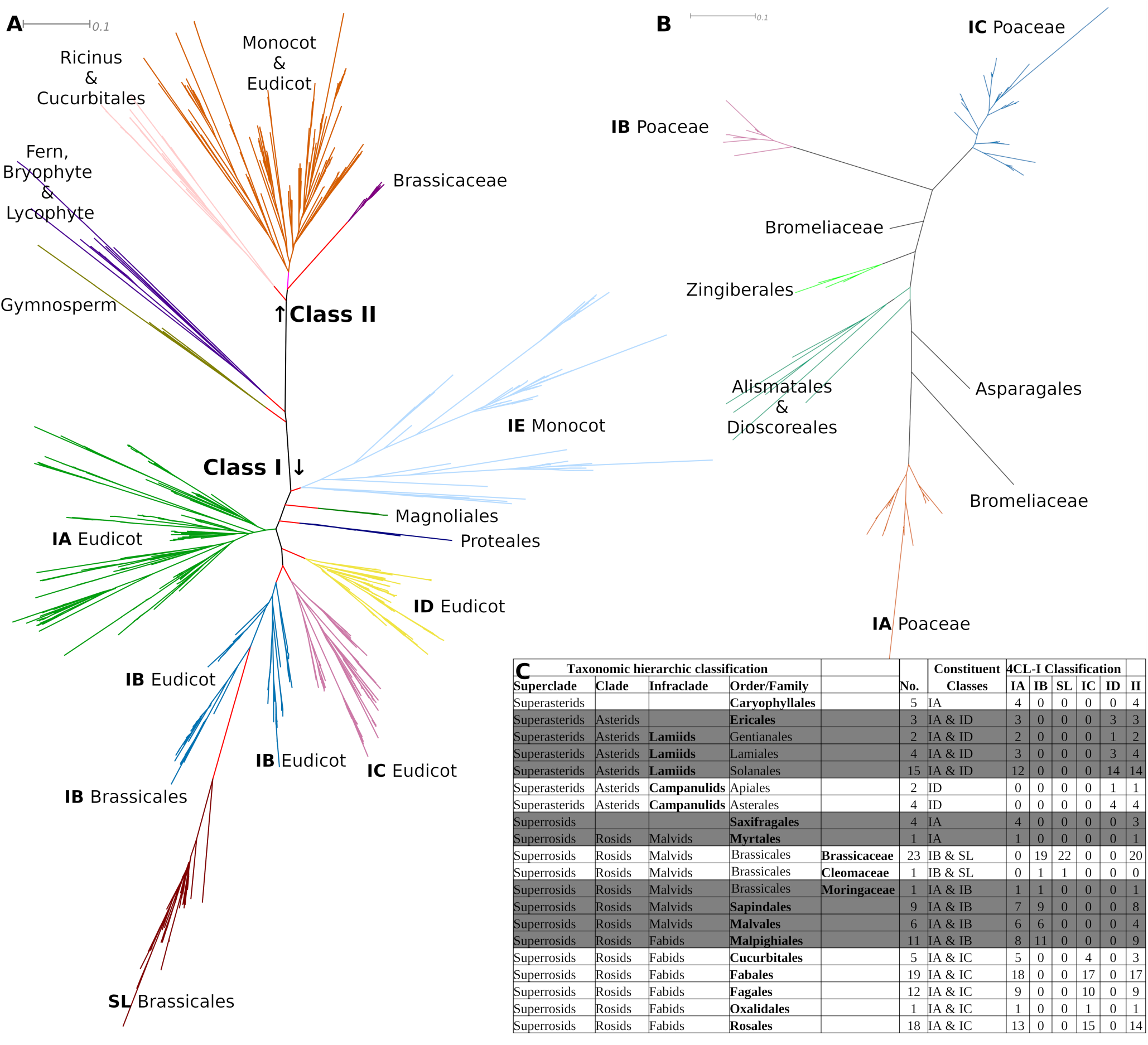
Phylogeny and Intuitive Clustering of Cytosolic 4CL Enzymes. For trees in A and B: Scale bars indicate the number of amino acid substitutions per site, reflecting evolutionary distances among taxa. Colours are randomly chosen to facilitate the identification of the selected clusters. Internal tree branches in A are in red to highlight the distances between the clades’ ancestral and basal nodes. **A subclustering of the Complete Cytosolic 4CL Family.** Numbers indicate statistical support of major bifurcations. Supplemental Document 3 has the bootstrapped tree in Newick format. Class I and II are separated by the presence of two non-angiosperm clades. Class I shows a separation of monocot (IE) and eudicot sequences (IA to ID) by two clades of early diverging angiosperms. Subclade IB contains the distant functional subfamilies of Sinapate:CoA Ligase (SL) (details in main text). **B Detail of Monocot Class I subclustering.** Poaceae Class I paralogues cluster into three subclasses. Non-Poaceae sequences occur in small intermediate clades. **C subclustering of Eudicot Class I Enzymes with Taxonomic Homologue Distribution.** The column headed with ‘No’. indicates the number of species from a certain order included in the analysis. ‘Constituent classes’ shows the subclasses that are represented in each of the orders. The subsequent columns show how many of the species of each order have at least one paralogue in each of the five indicated CI classes and in CII. Per species details are in Supplemental Document 2.

A first major finding is that the divergence of Class I and Class II occurred in the latest common angiosperm ancestor, before the divergence of monocots and eudicots. The cytosolic 4CL tree has two major, composite, clades representing Class I and II 4CL enzymes and containing only mono- and eudicot sequences. These large clades are topologically separated by two smaller clades. One contains all 4CL homologues from fern *A. filiculoides* (n=2), lycophyte *S. moellendorffii* (n=2) and bryophyte *Physcomitrella patens* (n=9), all at relatively large distances from their common ancestor. The other has all sequences from gymnosperms *P. taeda* and *Picea abies*, also at relatively large distances. This also clarifies that the paraphyletic Class II enzyme clade of figure 2, in fact constitutes a monophyletic group of angiosperm Class II homologues with a single gymnosperm sequence, which is now placed in a gymnosperm-specific 4CL clade. The apparent absence of canonical Class I and Class II homologues may correspond to the different lignin and suberin compositions found in conifers (Nakamura et al. 1974; Graça 2015). These nevertheless appear to have at least two homologues and may have undergone distinct diversification.

In correspondence to the phylogeny and classification of Figure 2, Class II enzymes showed only modest diversification giving rise to one major subclade and two minor subclades. Only part of this diversification resulted from redundancy since only *Ricinus* species have sequences in two of the subclades. A striking finding is that Class II enzymes from Brassicaceae, but not from other Brassicales, evolved separately under a strong functional constraint.

Class I homologues initially follow the species phylogeny, forming separate monophyletic clades for monocot and eudicot sequences. These are separated by two smaller clades representing the early-diverging angiosperm lineages Magnoliales and Proteales. Subsequently, the Class I homologues from Poaceae appear to have diversified into three subclasses, here designated Poaceae Class IA, IB and IC, with a single FH for most species included. This is best appreciated when looking at the incepted clade as midpoint-rooted tree (Figure 5b; *Repo-ClassIMonocot.nexml*). Smaller clades separate the Poaceae Class IA from Poaceae Classes IB and IC and contain sequences from non-Poaceae monocots. Species level details are found in Supplemental Document 2. We reviewed existing biochemical data for rice 4CLs (Gui et al. 2011) that show that Poaceae Class IA prefers ferulate and readily activates sinapate *in vitro*. Poaceae Class IB and Poaceae Class IC, the latter with two homologues in rice, do not activate sinapate and prefer coumarate or ferulate, while both Poaceae Class IC homologues are also active on caffeate.

Dicot Class I enzymes did not show clear signs of further diversification for the species from the orders of Caryophyllales, Saxifragales, and Myrtales, or the Campanulid infraclade. In the other orders redundancy, diversification, gene loss and possible hybridization events resulted in a topology that does not strictly follow taxonomy. Interestingly, species possess representatives from exactly two of the five eudicot CI subclasses we designated. Subclass IA is the most broadly distributed, with 14 out of the 17 analysed orders having at least one representative. This suggests it represents the ancestral Class I lineage in eudicots, substantiated by the fact that this is the eudicot Class I subclade that is most closely related to the monocot Class I subclade. By contrast, subclass ID is restricted to Asterids whereas subclass 1C is found in four out of five Fabid orders, while Malpighiales appear to have swapped subclass 1C for subclass 1B, based on evidence from all 11 Malpighiales species included (Supplemental Document 2).

Subclass IB and subclass SL form phylogenetic clade IB. The SL subclass we functionally annotated based on the well-described sinapate:CoA ligase from *A. thaliana.* This sequence (SwissProt identifier Q9LU36) was assigned as outlier and removed from the training set in order to allow automated, objective clustering. The sequence was identified by the eudicot Class I HMMER profile, together with SL orthologues from Brassicales species. They appear as a subfamily that more recently evolved in Brassicales only: All other Brassicales Class I homologues are found in the IB Brassicales subclade from which the SL subclade emerges (Figure 5a). The IB clade includes members from the other Malvid orders Sapindales and Malvales in other IB subclades. Besides this intricate pattern of diversification, it is striking that, with the exception of those orders represented by a single subclass, all species carry representatives from exactly two of the five designated subclasses, in the combinations of IA & IB, IA & IC, IA & ID, or IB & SL (Figure 5c).

### 4 Expression of the 4CL/ACS Gene Family in Potato

We analysed public RNA-seq data (Massa et al. 2011) using the eFP browser (Winter et al. 2007) (Supplemental Figure 3). St4CLIA is the major family transcript in mature tubers, particularly in peel, and is expressed to some extent in stem. St4CLID1, ID2 and ID3 are active in stem and petiole, with St4CLID1 and ID2 also active in tuber. This can be explained as being involved in developmental suberin and lignin production. St4CLII, with transcripts in Phureja but not Tuberosum tuber, and St4CLIII, transcripts mostly in flower, do not appear as directly related to their suggested functions. Homologues StOPCL, StUACSI and StUACSII2 have little or no expression in tubers but are expressed in most other organs, in correspondence with a role for instance in the JA pathway. These show significant differences between the Phureja and Tuberosum cultivars. No RNA-seq data for StACOS, StUACSII1 or StUACSIII were found.

In order to confirm the classification of potato cytosolic 4CL genes, we analysed St4CL transcript levels in Andean potato tubers, accompanied by the levels of StCHS and StFHT as markers for flavonoid metabolism and suberin/lignin production, respectively. In wound periderm, the anthocyanin pathway has been shown to be strongly downregulated whereas suberin related genes were upregulated (Fogelman et al. 2015). Therefore, tubers from the red flesh and skin Santa María variety were subjected to wounding and transcript levels of the five St4CLs genes were determined by qRT-PCR (Figure 6). Peel samples (native periderm) were included as control, note that these effectively represent the peel whereas wound periderm samples are represented by 2-3mm cut tuber slices. St4CLID1 and St4CLID2 as well as StFHT expression was induced by wounding, whereas St4CLII and StCHS were repressed (Figure 6). Consistently, levels of anthocyanins content diminished in wounded Santa María tubers (Figure 6).The levels of expression of St4CLID2 were three orders of magnitude lower than St4CLID1, whereas in the public dataset these showed similar transcript levels. St4CLIA and St4CLID3 transcripts were not detected in native or wounded periderm and showed the lowest absolute levels of all Class I genes (Supplemental Figure 3).

**Fig 6.**
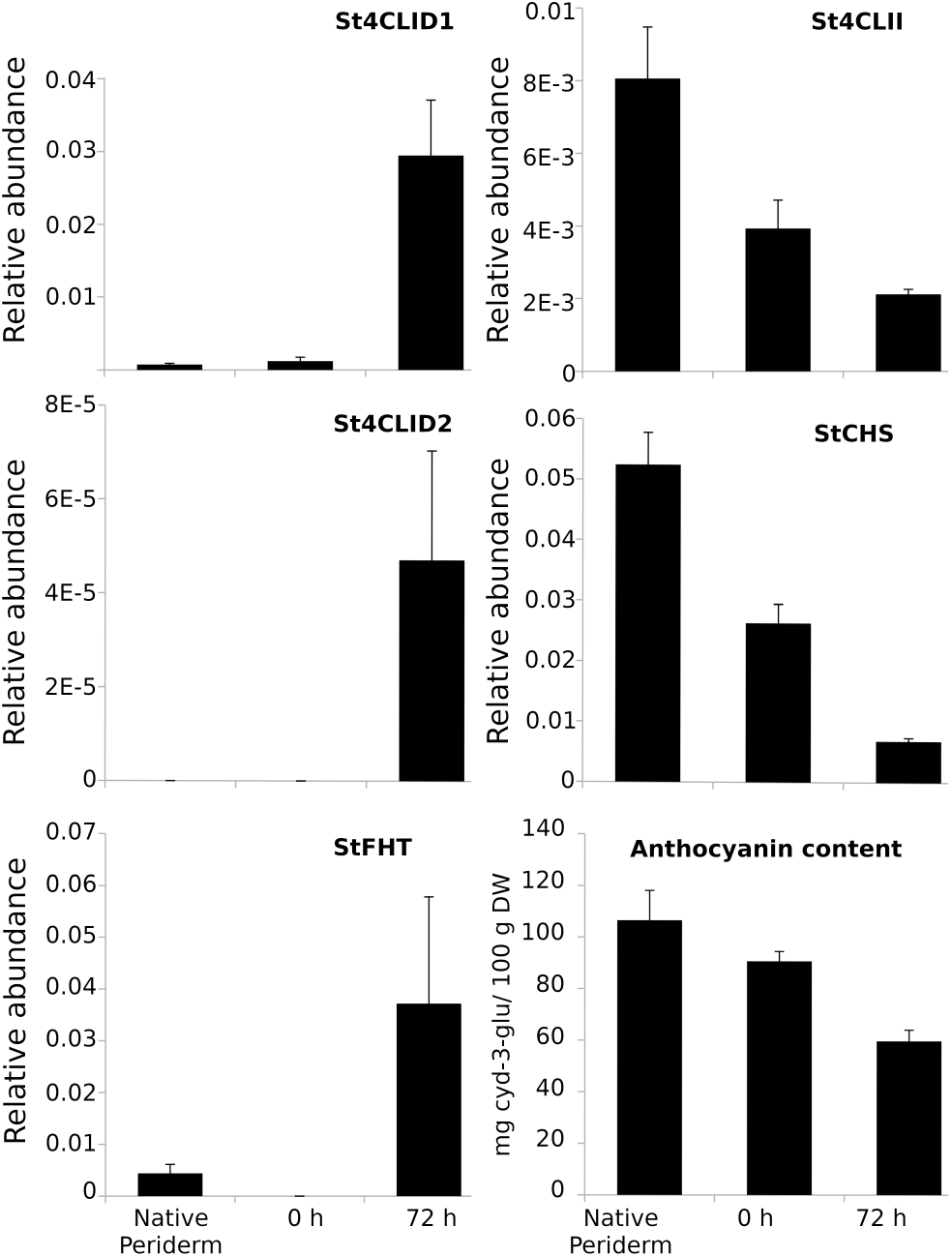
Expression profile of St4CL genes and anthocyanin content in potato tubers subjected to wounding. Transcript levels of St4CL genes were analyzed by qRT-PCR in wounded Santa María potato tubers. Data represent mean ± SE values from two independent experiments.

Next we studied four Andean potato varieties with contrasting flesh/skin tuber colours (Table 1). As it can be seen in Figure 7, Santa María variety, which is intensely pigmented in flesh and skin, showed the highest expression levels of both StCHS and St4CLII, skin samples showing levels 2 to 4 times higher than flesh. Chaqueña and Waicha varieties showed low levels but with the same pattern when comparing skin and flesh. Moradita showed surprisingly high levels of St4CLII. Transcript levels of St4CLID1 were extremely low and appeared as constitutive (Figure 7). St4CLIA and St4CLID2 and ID3 transcripts were not detected in flesh nor skin of any of the potato cultivars. StFHT was consistently detected in skin but not in flesh. All together, results show that St4CLID1 and St4CLID2 transcript levels increase upon wounding, in correspondence with an induction of monolignol and suberin biosynthesis. Similarly, St4CLII appears to be involved in flavonoid biosynthesis, confirming it as a Class II enzyme.

**Fig 7.**
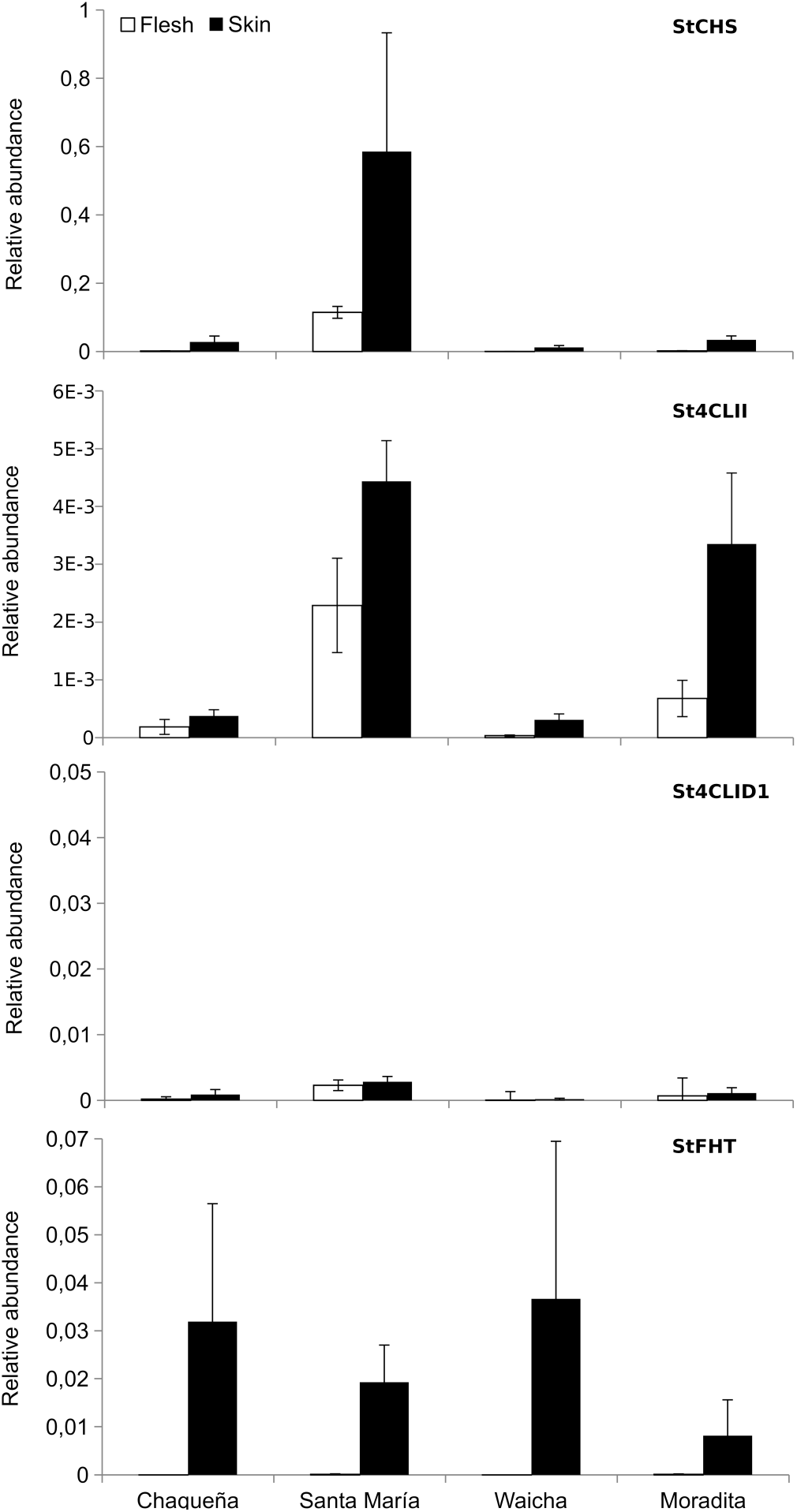
Expression profile of St4CL genes in flesh and skin of tubers from Andean potato varieties. qRT-PCR data represent mean ± SE values from three independent biological samples. The vertical (Y-axis) ranges for all genes is set similar to that in Figure 6, allowing for direct visual comparison of data magnitude.

## Discussion

We study the biosynthesis of phenylpropanoids, anthocyanins and other flavonoids in potato tuber. Previously, we showed an important role for CGA as a central compound in the phenylpropanoid metabolism, related to both suberin deposition and anthocyanin production in Andean potato tubers (Valiñas et al. 2015, 2017). Here we set out to study the role of 4CL. In order to be able to identify all possibly involved 4CL homologues, we decided to classify the corresponding superfamily. We performed a profound analysis given an inconsistent description of the gene family in literature; the existence of outdated SwissProt annotation lines; and, most importantly, the hierarchic nature of protein superfamilies and the corresponding problem of precision and recall. We combined various approaches and software for automated sequence scrutiny and sequence clustering to achieve high P&R on a dataset with both annotated, curated sequences from SwissProt, as well as genome-wide dataset from 15 plant species. The sequences were classified into eight clear functional subfamilies. The largest of these subfamilies contains the Class I 4CL enzymes that is under neo-functionalisation, which required a sequence set of 215 plant species for classification into various functional subclasses.

Since protein families are hierarchic in nature, we performed a more sensitive sequence mining in parallel. A BLAST search using the consensus E-value cut-off of 1E-5 rather than the strict 1E-50 resulted in a larger initial set of sequences and, correspondingly, a more sensitive HMMER profile that identified an approximately double-sized set of sequences (Supplemental Figure 4). Interestingly, searching the corresponding tree with the terms 4CL and 4CL-like, identified a clade with the exact same sequences as identified in the strict approach. Thus, the strict sequence mining shows excellent P&R and includes all sequences annotated as 4CL or 4CL-like. The definition of the superfamily is also defined by the need to include Class III 4CL enzymes. The sequence mining of the 200 additional plant species was performed using HMMERCTTER classification (Pagnuco et al. 2018), which is based on 100% P&R-SD, and taxonomic contribution analyses showed no inconsistencies.

The representative sequence set not only allowed an easy identification of cytosolic 4CL sequences from potato, the phylogeny-based classification was, besides the cytosolic 4CL clade with the Class II family and the complex Class I family, surprisingly simple. The six other clades have relatively large distances between the ancestral and basal nodes, and mostly a single FH per species, thereby corresponding to functional subfamilies. A possible exception is UACSII that shows 2 or 3 FH for certain species and possible subclades (Figure 1). Since there is no known function of UACSII sequences we did not further subcluster this clade. We designated three subfamilies hitherto annotated as 4CL-like as ACOS5, OPCL and Class III 4CL using data from the literature (Koo et al. 2006; Kienow et al. 2008; De Azevedo Souza et al. 2009; Block et al. 2014). The data-mining of the additional 200 plant species showed similar taxonomic trends as the that of the representative sequence set but, with a higher resolution. The UACSIII clade lacks Poaceae and Brassicaceae sequences, rather than monocot and Arabidopis. Numerical analysis shows it lacks many species. UACSII, UACSI, OPCL, 4CL Class III, ACOS5 and UACSIII lack homologues in 18, 16, 25, 33, 24 and 45 species, respectively, from a set of 136 eudicot species excluding the Brassicales (Supplemental Document 2). This apparent lack of UACSIII homologues suggests its function is not crucial.

We studied the cytosolic 4CL classes more profoundly since most species contain more than one Class I FH. Besides that the cytosolic 4CL tree (Figure 5a) points out that the diversification of Class I and II occurred in the common ancestor of angiosperms, it also shows that Class I enzymes are likely in a process of functional redundancy and diversification. Not only are Class I enzymes represented by more than a single FH, the clade show clustering that points out different subfamilies. Unfortunately, their classification is obscured by the fact that phylogenetic trees reflect drift and shift. Most of the major clades representing functional families in figure 2 have monocot and eudicot sequences in different subclades, (*Repo-4CLACS_Representative.nexml*) albeit the intertwined signals from functional and taxonomic diversification, could be easily disentangled. This was hampered for Class I sequences, since these sequences have evolved more recently, demonstrated by a taxonomic grouping that occurs at hierarchically lower levels.

We hypothesise a chain of evolutionary events that starts with the latest common angiosperm ancestor that had a single 4CL Class I enzyme. Upon the bifurcation into monocot and eudicot, these first separated by means of drift that was followed by various, independent duplications. In the ancestor of the Poaceae, two subsequent duplications occurred, leading to three subclasses under a different functional constraint (Figure 5b). In eudicots, duplications occurred later in the hierarchic taxonomy, hence, evolution led to a more complex clustering pattern (Figure 5c), most species having Class I paralogues in two of five Class I subfamilies. Eudicot class IA was the ancestral eudicot Class I clade (Figure 5a) and we assume that this ancestral Class I family did show high substrate permissiveness with low activity with the voluminous sinapate. The latest common ancestor of Asterids showed a single duplication, resulting in the subfamily designated ID. In Campanulids, the original IA homologue got lost. Rosids showed different duplications in different lineages, somewhat obscured by death and possibly hybridisation events. In Malvids this initially led to the novel IB subfamily. Next, in Brassicaceae (and Cleomaceae), the IB homologue duplicated and one of the duplicates underwent a deletion allowing sinapate:CoA ligase activity. Brassicaceae IB evolved towards simple substrate as cinnamate and caffeate and the existence of the novel sinapate:CoA ligase allowed for the loss of the original, permissive IA homologue. Other Malvid orders maintained the IA besides the IB homologue. The presence of IB in the Fabid order of Malpighiales might be explained by hybridisation or a horizontal transfer. The remaining Fabid orders evolved the IC subfamily. There are no data to support any prediction on substrate preference for IC and ID subfamilies. We merely assume they are more specific than IA.

We did not compare different Class I subfamilies by further computational analysis. This was mostly motivated by the results from the eight family comparisons of the hallmark subsequences of Box I and Box II (Figure 3), and the predicted binding pocket. The boxes are relevant as hallmarks for, respectively, adenylating enzymes and cytosolic, but not peroxisomal, 4CL sequences. They are not informative when it concerns substrate specificity predictions. The predicted binding pocket appears as a better element to make substrate related predictions as is seen by the high number of substitutions shown between cytosolic 4CL enzymes and ACOS5. Few differences were found in the comparison of Class I and Class II binding pockets; with Y239F and Y442F being the most promising substitutions. Position 239 is found at the interface of both binding pockets. Mutational analysis performed by Li and Nair (Li and Nair 2015) showed that this did affect K_m_ towards coumarate. Y442F concerns a position in the CoA pocket and the difference given by a single hydroxyl can be envisaged to affect CoA binding, favouring binding to Class I above Class II enzymes. We suggest CoA concentration rather than hydroxycinnamate concentration regulates cytosolic 4CL activity and as such downstream fluxes. With both enzymes present and low CoA concentrations, only Class I is active, probably also acting on p-coumarate (Figure 1). When energetically more favourable conditions occur, CoA concentrations likely increase thereby activating Class II enzymes and shifting metabolite flux towards flavonoids.

The transcript analyses of the predicted Class I and Class II 4CL in potato substantiate their classification, albeit that analyses of public data were not overly informative. Comparison of tissues that contained different levels of anthocyanins and particularly the wound response experiment were more insightful. Wounding leads to an increase of Class I transcript thereby shifting the metabolic flux towards lignin and suberin deposition. Interestingly, both St4CLID homologues are induced but not the ST4CLIA homologue. This provides a feasible, alternative explanation for the evolution of Class I enzymes: two different classes have evolved in order to more efficiently deal with developmental and stress-related requirements of lignin and suberin production. This would not necessarily oppose a substrate driven diversification since developmental and wound related lignin and suberin may have different proportions of hydroxycinnamate derived moieties. Profound chemical analysis of native and wound periderm may as such be more informative than dedicated enzymatic assays *in vitro.* Both lie beyond the scope of the present work and our main interest. Enzymes assays may be informative to investigate the hypothesised role of CoA concentration.

## Data repositions

All sequence data used for this work are public. In order to improve transparency and reproducibility of our research we have made all important datasets (fasta files with sequences, aligned sequences, HMMER profiles and nexml-formatted treefiles of bootstrapped trees) available via the CONICET data repository at https://ri.conicet.gov.ar/handle/11336/290620. These datafiles are indicated with *Repo-filename* throughout the Results section.

