## Supplemental Table 1 for "Classification and Evolution of the Plant 4CL/ACS Superfamily and Transcript Analysis in Andean Potato"

**Supplemental Table 1: Primer sequences used for qRT-PCR**

| Gene | Primer | Sequence (5’ to 3’) |
|---|---|---|
| St4CLID3 | fw | GGATGAAATCAAGGATTTTATCTCT |
| | rv | CCTGATGGAGATTTAGGAATCA |
| St4CLIA | fw | ATTTGTAGTGAAAGCAAATGGTAG |
| | rv | CTGATGGAGCTTTTGGAATTTCT |
| St4CLID1 | fw | TTGTTAGATCAAATGGCTCCACA |
| | rv | CCCGATGGAGATTTAGGAACA |
| St4CLID2 | fw | AGAATGATAGATGAACAAGCG |
| | rv | CGGAGATTTTGGTACCGTC |
| St4CLII | fw | TCGGCCCAAGGTTTTGATC |
| | rv | AGACTTCGGAATCGCGTGA |
| StFHT | fw | GCCTGATCCTGTTACACTTGG |
| | rv | AAACATGCAATGGTTCATGC |
| StCHS | fw | CCGCTATCATTATGGGTTCG |
| | rv | GGAACATCCTTGAGTAAGTGGAAC |
| StL2 | fw | CGAAGGAGCTGTTGTTTGTAAC |
| | rv | GGGCACAATCTTTTTGGC |
