## Supplemental Table 2 for "Classification and Evolution of the Plant 4CL/ACS Superfamily and Transcript Analysis in Andean Potato"

***Supplemental Table 2: Binding pocket definitions of Nt4CL2 3D structures co-crystallized with various substrate analogues.*** *Highlighted positions are within 5 Å of the indicated substrate analogue. The ATP, AMP, and hydroxycinnamoyl moieties share the same pocket (yellow highlight), while CoA binds in a distinct but adjacent pocket (red highlight). Note that 5BSR contains both an AMP and a CoA moiety. The P loop is part of Box I; Box II does not correspond to a defined binding pocket. Pockets were determined using PyMOL [52].*

| | P loop | | | | | | | | A7 | | | | | |
|---|---|---|---|---|---|---|---|---|---|---|---|---|---|---|
| | 189-197 | 213 | 232-243 | 260-261 | 281-284 | 306-312 | 331-344 | 360 | 418-420 | 432-446 | 506 | 509 | 524 | 526 |
| 5BSM ATP | | | | | | | | | | | | | | |
| 5BSR-AMP | | | | | | | | | | | | | | |
| 5BST Cou-AMP | | | | | | | | | | | | | | |
| 5BSU Caf-AMP | | | | | | | | | | | | | | |
| 5BSV Fer-AMP | | | | | | | | | | | | | | |
| 5BSR-CoA | | | | | | | | | | | | | | |
