## Supplementary material for "Classification and Evolution of the Plant 4CL/ACS Superfamily and Transcript Analysis in Andean Potato": Supplemantal Figure 1

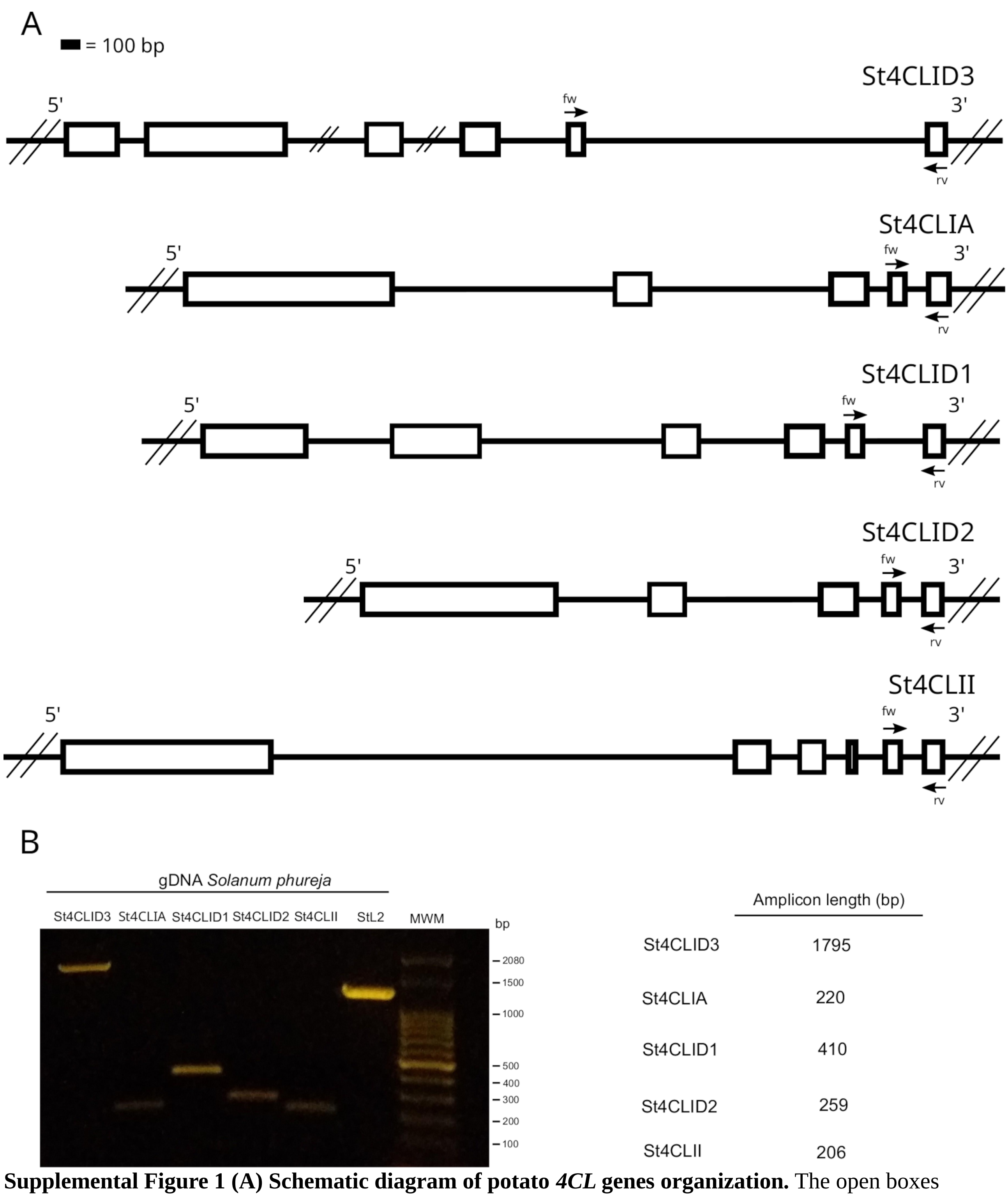


**Supplemental Figure 1 (A) Schematic diagram of potato *4CL* genes organization.** The open boxes represent coding sequence. The locations where primers anneal are shown. **(B)** PCR products of isoform-specific primers using as a template gDNA from *Solanum phureja*.
