## Supplemental Figure 2 for "Classification and Evolution of the Plant 4CL/ACS Superfamily and Transcript Analysis in Andean Potato"

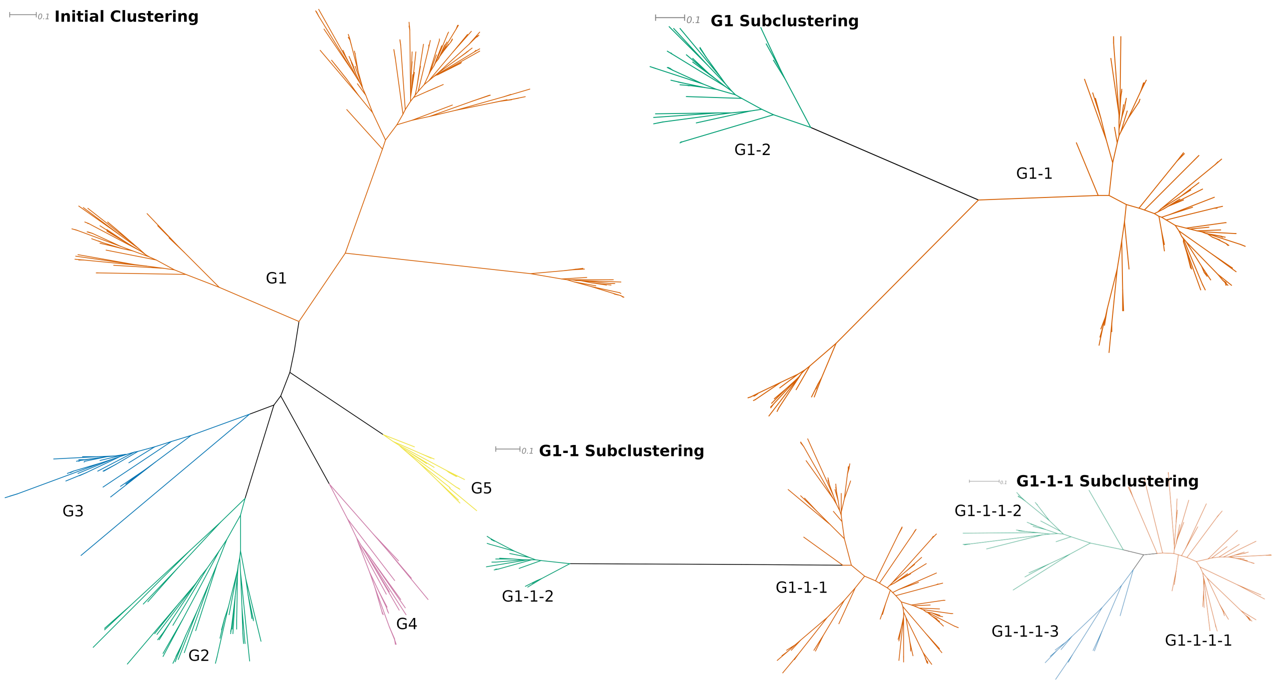


***Supplemental Figure 1: Hierarchic clustering of the representative 4CL/ACS sequence set.*** *HMMERCTTER identifies the largest clades that are 100% P&R-SD.* ***Initial clustering:*** *Cluster G2 to G5 are well separated and have single functionally non-redundant orthologues (See Table 1 main document) and were accepted as final clusters. The G1 cluster has at least three subclusters, based on the same criteria. The clade was extracted from the complete tree and reclustered with HMMERCTTER.* ***G1 Subclustering:*** *G1-2 is a functional subcluster; G1-1 is reclustered.* ***G1-1 Subclustering:*** *G-1-1-2 is is a functional subcluster; G1-1-1 is reclustered.* ***G1-1-1 Subclustering:*** *G1-1-1-1 and G1-1-1-3 are the Class I subcluster separated by HMMERCTTER into taxonomic subclusters. G1-1-1-2 is the Class II subcluster. Scale bars show the number of amino acid substitutions per site reflecting evolutionary distances of the different taxa.*

Supplemental Figure 2: Classification and Evolution of the Plant 4CL/ACS Superfamily and Transcript Analysis in Andean Potato. Valiñas, Lanteri, Andreu and ten Have
