## Supplemental Figure 3 for "Classification and Evolution of the Plant 4CL/ACS Superfamily and Transcript Analysis in Andean Potato"

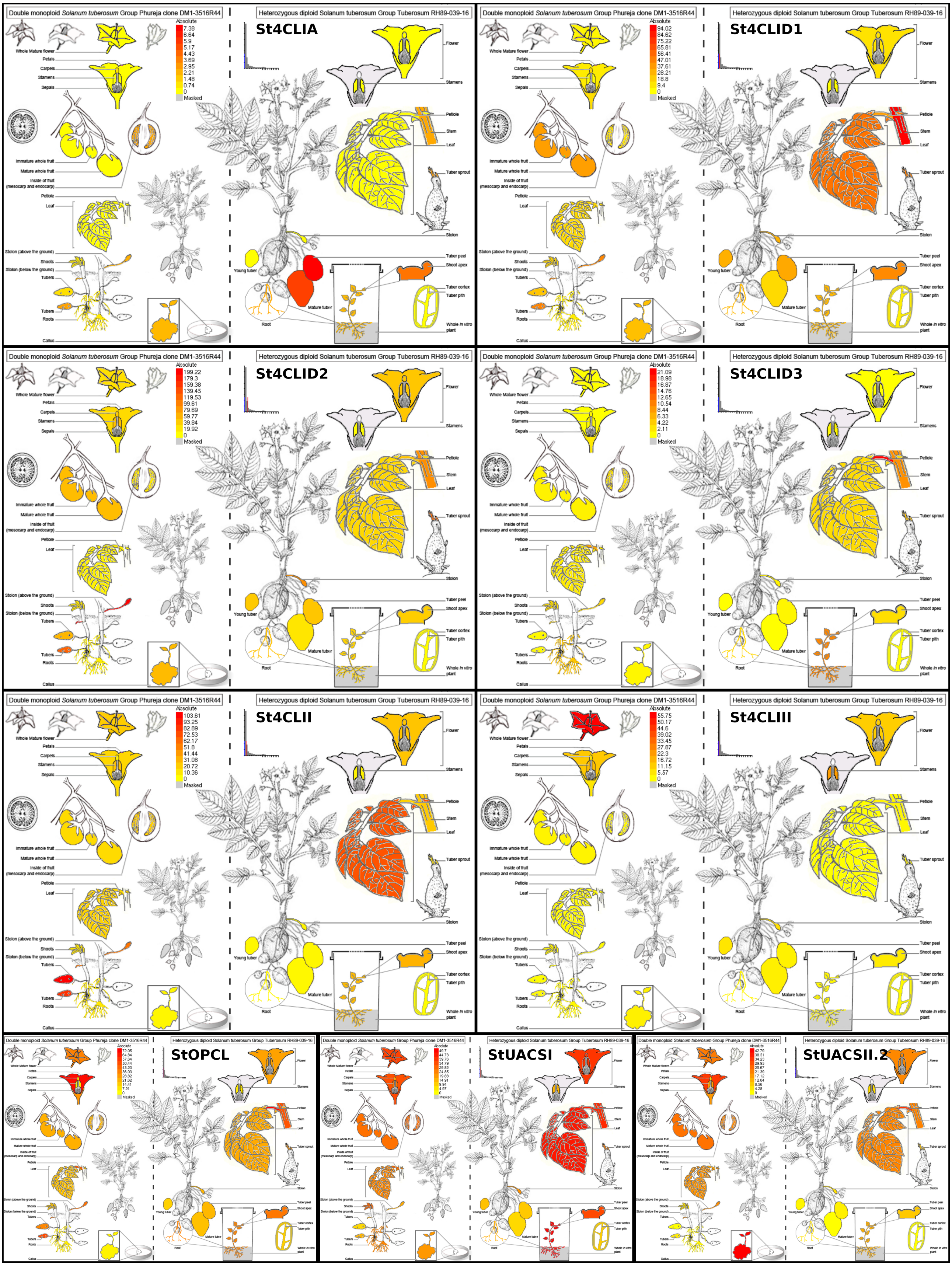


**Supplemental Figure 3: eFP browser view of tissue and developmental gene expression 4CL/ACS gene family.** Each panel shows the expression in double monoploid *S. tuberosum* group *phureja* (left) and in the heterozygous diploid *S. tuberosum* group *tuberosum* (right). The heatmap shows relative expression levels, not that yellow corresponds to no transcripts detected. No data were found for StACOS, StUACSI and StUACSIII.

Supplemental Figure 3: Classification and Evolution of the Plant 4CL/ACS Superfamily and Transcript Analysis in Andean Potato. Valiñas, Lanteri, Andreu and ten Have
