## Supplemental Figure 4 for "Classification and Evolution of the Plant 4CL/ACS Superfamily and Transcript Analysis in Andean Potato"

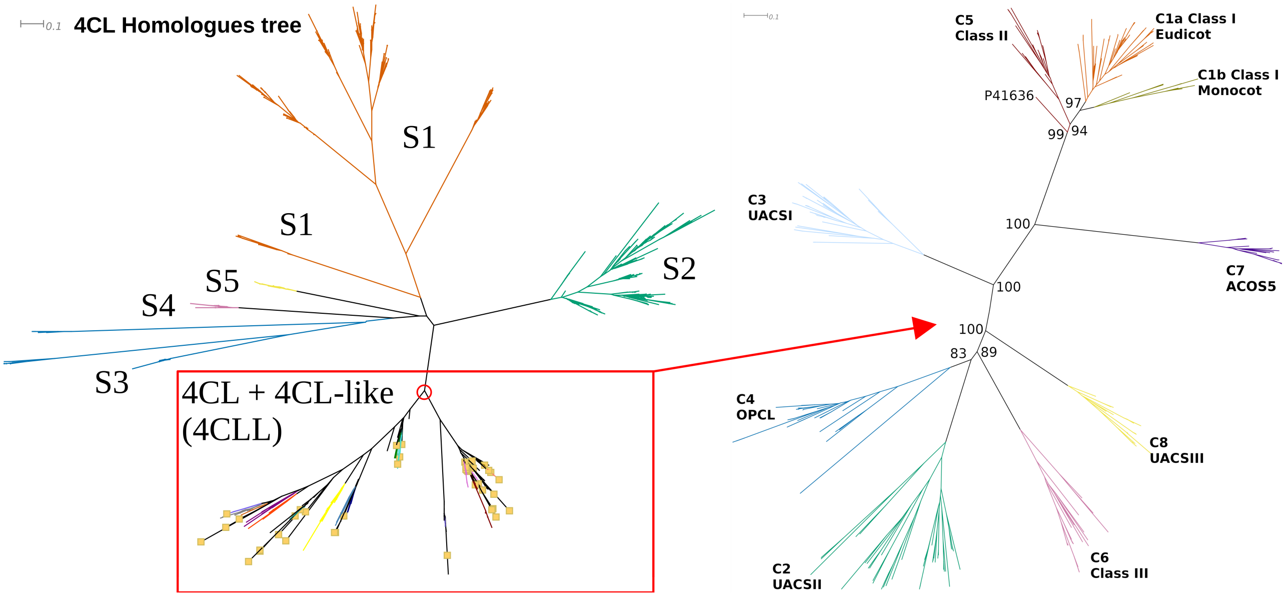


***Supplemental Figure 4: The 4CL/ACS family is actually a subfamily of the superfamily of long-chain-fatty-acid-CoA ligases.*** *The 4CL homologues tree has all sequences obtained using a more sensitive BLAST and corresponding hmmsearch. Besides the 4CL/ACS family (not 100% P&R-SD and with shattered clustering due to the presence of outlier* Q9LU36 (SL)), *we identified 5 additional 100% P&R-SD clades (S1 to S5) representing other subfamilies of the long-chain-fatty-acid-CoA ligase superfamily. All sequences that contain CL (hence including 4CLL) in their assignation line are selected by a beige square, all fall within the 4CL/ACS clade (red box) that was identical to our 226 sequence set we obtained as described in the main document.*
